# Potent optogenetic inhibition of behavior with anion channelrhodopsins

**DOI:** 10.1101/082255

**Authors:** Farhan Mohammad, James C. Stewart, Stanislav Ott, Katarina Chlebikova, Jia Yi Chua, Tong-Wey Koh, Joses Ho, Adam Claridge-Chang

## Abstract

Optogenetics employs light exposure to manipulate physiology in genetically modified organisms. There are abundant tools for optogenetic excitation of neuronal activity, but the limitations of current activity photo-inhibitors present an obstacle to demonstrating the necessity of specific neuronal circuits. Here we show that anion channelrhodopsins can be used to specifically and rapidly inhibit a range of systems involved in *Drosophila* locomotion, wing expansion, memory retrieval and gustation, demonstrating their broad utility to the circuit analysis of behavior.

## Introduction

Our ability to understand the neuronal control of behavior has been transformed by the advent of techniques enabling researchers to precisely manipulate neuronal activity in transgenic animals (Sweeney et al. 1995; Johns et al. 1999; Kitamoto 2001; Zemelman et al. 2002; Boyden et al. 2005; Roth 2016; Hamada et al. 2008; Tye and Deisseroth 2012). Optogenetics is one such technique which uses controlled light exposure to reversibly modulate the activity of light-sensitive ion channels. Optogenetic actuators can be expressed in genetically–modified organisms with circuit specificity (Zemelman et al. 2002) in behaving animals (Lima and Miesenböck 2005) with millisecond–resolution (Boyden et al. 2005). However, while optogenetic activators are in widespread use, the toolkit of optogenetic inhibitors of neuronal activity is more limited: consequently, in a number of experimental systems it is relatively much easier to show that a specific neural circuit is sufficient for any given function, yet demonstrating the necessity for that circuit remains problematic. Suppression of neuronal excitability uses either an influx of anions or an efflux of cations; this has previously been achieved using light-driven chloride (Zhang et al. 2007) or proton pumps (Chow et al. 2010), but they require high-density expression and intense light for effective inhibition, limiting their utility. Recently, a new class of inhibitory, anion-conducting, light-gated channels has been developed, including channelrhodopsins engineered to conduct chloride ions (Wietek et al. 2014; Berndt et al. 2014), and two naturally-evolved anion channelrhodopsins cloned from an alga (Govorunova et al. 2015). The algal *Guillardia theta* anion channelrhodopsins (GtACRs) possess several attractive features as optogenetic inhibitors: they have much higher conductances than other inhibitory optogenetic tools, are rapidly responsive, require only low light intensities for activation, and are comprised of both a cyan-gated (GtACR1, maximal sensitivity at 515 nm) and a blue-gated channel (GtACR2, maximal sensitivity at 470 nm) (Govorunova et al. 2015). Considering these properties, we hypothesized that the GtACRs could be employed to effectively optogenetically inhibit neuronal circuits in a model organism. Here we show that both GtACR1 and GtACR2 potently, rapidly, and reversibly inhibit a range of behaviors in *Drosophila* during controlled exposure to consumer light sources.

## Results

### Climbing flies fall when GtACRs are light-activated in cholinergic neurons

Genes coding for the GtACRs were expressed in cells that release the neurotransmitter acetylcholine (Kitamoto 2001; Salvaterra and Kitamoto 2001). While climbing on a vertical surface, *Drosophila* expressing the cyan-gated GtACR1 fell when exposed to just 1.3 µW/mm^2^ green light from a projector (Figure 1A, Video 1). GtACR1 flies (*Cha>GtACR1*) were also susceptible to blue and red light (at ≥14 µW/mm^2^ and 39 µW/mm^2^ respectively). Flies expressing the blue-gated GtACR2 in cholinergic neurons fell in response to blue light only at 14 µW/mm^2^ (Figure 1B) and to both blue and green light at the highest intensities (but not red light). By contrast, flies expressing enhanced *Natronomonas pharaonis* halorhodopsin (eNpHR) did not fall when illuminated with 39 µW/mm^2^ red projector light. Control flies were unaffected by light (Figure 2A).

**Figure 1:**
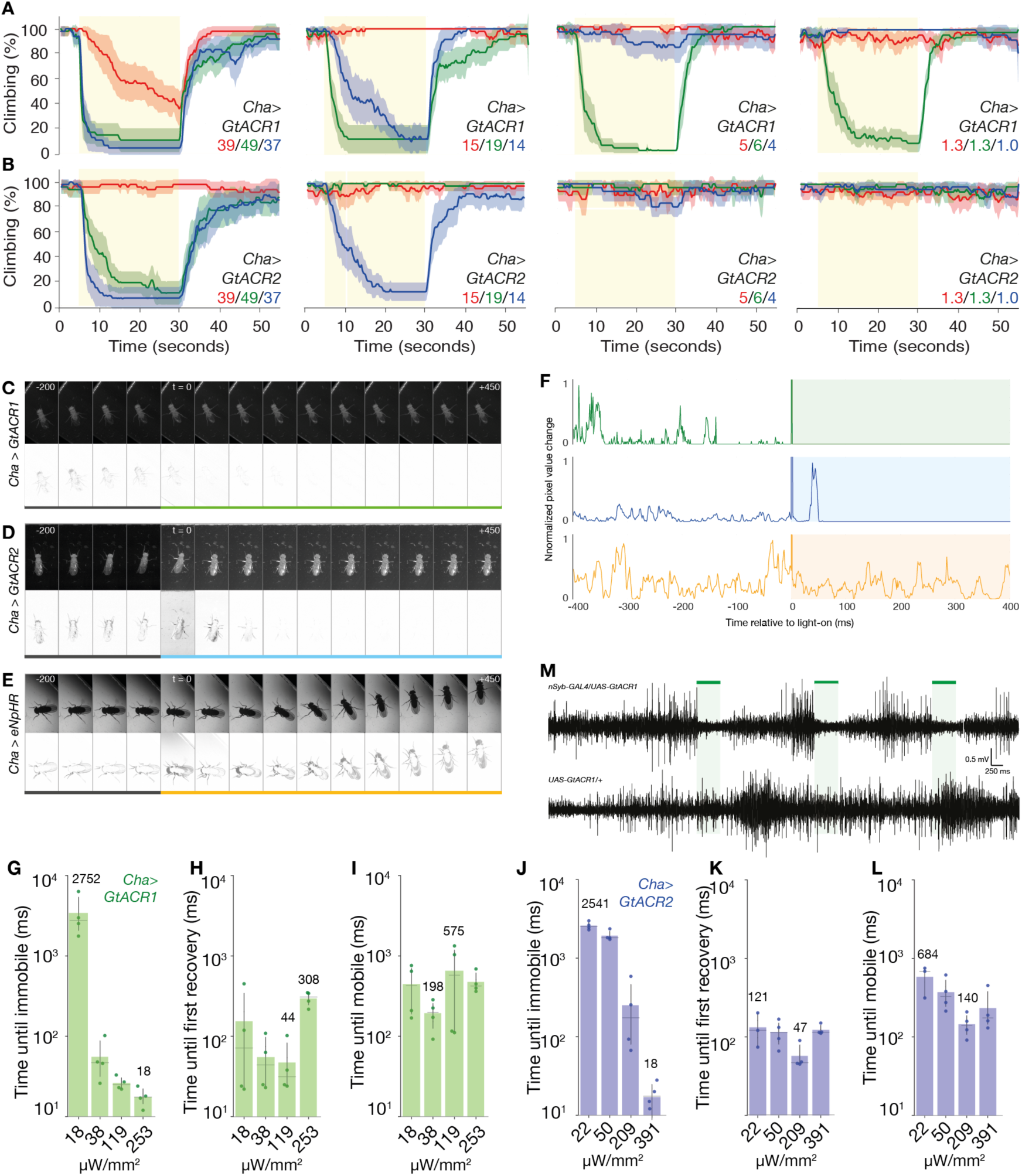
*Guillardia theta* anion channelrhodopsins are potent inhibitors of motor function and neuronal spiking. **A.** Flies expressing GtACR1 in cholinergic neurons (*Cha-Gal4>UAS-GtACR1*) fell from a vertical surface when illuminated. Light intensities in μW/mm^2^ are given in each panel. The proportion of climbing flies is given as a percentage of flies outside the floor area of the chamber. At the highest setting, flies fell in response to green, blue and red light, but responded only to green light at lower intensities. Each solid trace is the mean of 3 experiments (48 flies), line color corresponds to the color of illumination from a projector, error ribbons are 95% confidence intervals. **B.** Flies expressing GtACR2 (*Cha>GtACR2*) fell in response to blue and green light at the highest intensity; green light had little effect at 19 μW/mm^2^, blue light (14 μW/mm^2^) was sufficient to causing falling. **C.** Representative Muybridge series from a high-speed video (1000 frames per second) during light onset (top row) show that green light illumination (peak 525 nm, 38 μW/mm^2^) of a fly expressing GtACR1 in cholinergic neurons elicited a rapid onset of immobility. Frames are spaced at 50 ms intervals, numbers indicate time relative to illumination start in ms. Difference images between consecutive frames (bottom row) map the motion. Coloured bar indicates frames where the fly is illuminated. **D.** Muybridge series indicates that illumination with 253 μW/mm^2^ of blue light rapidly paralyzed the *Cha-Gal4>UAS-GtACR2* subject fly. Upper row shows video frames, lower row shows difference images. **E.** A representative series shows that illumination with amber light (peak 591 nm, 495 μW/mm^2^) failed to interrupt the movement of a *Cha-Gal4>UAS-eNpHR* fly. **F.** Difference images from the videos in panels C-E were used to quantify motion. Plots show the normalized sum of pixel value differences across a region of interest that contained the moving fly. *Cha-Gal4>* GtACR flies were motionless within 100 ms, while illuminated *Cha-Gal4>*eNpHR flies showed similar pixel changes before and during light exposure. The pixel difference spike observed at light onset is due to imperfect exclusion of visible light by the longpass filter. **G.** Four *Cha-Gal4>UAS-GtACR1* flies were recorded at 1000 fps and frame-by-frame inspection was used to quantify the time until each fly was immobilized after lights-on. The horizontal axis indicates the four light intensities tested, the numerals in the panel indicate the medians of the fastest and slowest responses. The time until immobile shortened dramatically with increasing green light intensity (peak 525 nm). Error bars are 95% confidence intervals. **H-I.** Times between cessation of illumination and the first sign of recovery, and times to full mobility, over four light intensities. There was no strong relationship between light intensity and recovery times. **J-L.** Times of immobility onset and recovery of four *Cha-Gal4>UAS-GtACR2* flies over a range of intensities. **M.** Representative recordings from larval segmental nerves. Pulses of green light (500 ms, 24 μW/mm^2^, green lines and shading) inhibited action potentials in nerves expressing GtACR1 (*nSyb>GtACR1,* pan-neuronal expression), but not in control larvae (*UAS-GtACR1*/+).

**Figure 2.**
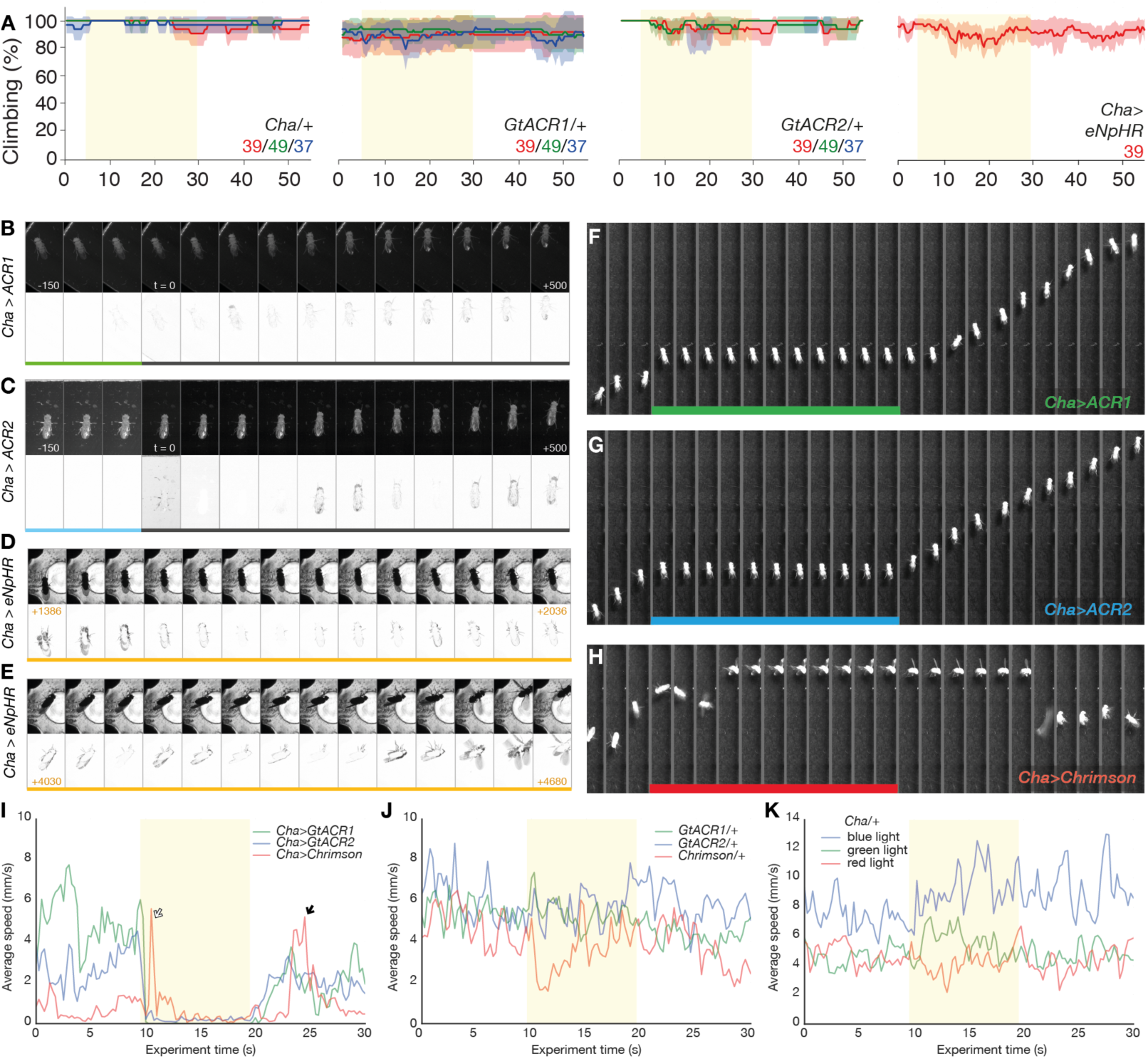
Anion channelrhodopsin action in the *Drosophila* cholinergic system. **A.** Control flies carrying either the driver or GtACR transgenes alone did not fall when illuminated with different wavelengths and intensities of light. The indicated colored light intensities are shown in μW/mm^2^. Similarly, the climbing performance of *Drosophila* expressing eNpHR in cholinergic cells remained largely unimpaired when the flies were illuminated with 39 μW/mm^2^ of red light. **B.** Muybridge series illustrates the recovery of a *Cha-Gal4>UAS-GtACR1* fly after light-off. Motion in the difference series prior to lights-off indicates that the fly was immobile, though not motionless. Inter-frame steps are 50 ms. **C.** Recovery of a *Cha-Gal4>UAS-GtACR2* fly over a 500 ms interval after light-off. **D–E.** A fly with a *Cha-Gal4>UAS-eNpHR* genotype exposed to 1900 μW/mm^2^ (i.e. sitting 3 mm above the LED) was only sporadically immobilized. **F.** A *Cha>GtACR1* fly entered into a static paralysis upon illumination with green projector light (92 μW/mm^2^). Frames are separated by 1 s intervals. **G.** A *Cha>GtACR2* fly underwent static immobilization during exposure to blue light (67 μW/mm^2^), though retained some leg movement. **H.** A *Cha>CsChrimson* fly underwent an active convulsion for several seconds upon illumination with red light (70 μW/mm^2^). During paralysis, the wings were extended; after light-off, seizure continued for ~6 seconds and was followed by another active convulsion before the fly regained a standing pose (last frame). **I.** Average speed of the three strains illustrated in F-H, before during and after exposure to projector light (*Cha>GtACR1,* green 92 μW/mm^2^; *Cha>GtACR2*, blue 67 μW/mm^2^; *Cha>CsChrimson*, red 70 μW/mm^2^). Lines are the mean speed of seven flies of each genotype. The GtACR flies are rapidly immobilized; the CsChrimson flies had increased speed after light-on that is related to convulsions (white arrow). The *Cha>CsChrimson* flies also convulse during recovery (black arrow). **J.** The walking speed of responder control flies is largely unchanged by light exposure, N = 15, 15, 15. **K.** The driver control (*Cha-Gal4/+*) line is unresponsive to projector illumination of any color, N = 15, 15, 15.

### GtACR activation in cholinergic neurons induces rapid, complete and reversible paralysis

Some of the fallen *Drosophila* moved during GtACR activation (Video 1), revealing incomplete inhibition of Cha^+^ neurons. Examining the motion of individual flies with a high frame-rate camera revealed that illumination with green light using light-emitting diodes (LEDs) induced complete and rapid immobility in *Cha>GtACR1* flies with intensities 38 µW/mm^2^ and above (Figure 1C). At higher powers, immobilization was observed in a fraction of all-*trans*-retinal-fed *UAS-GtACR1*/+ flies (36% [95CI 12, 68] at 253 µW/mm^2^, N = 11; 13% [95CI 2, 31] at 119 µW/mm^2^, N = 15; zero at the lower intensities, N = 12, 9), possibly due to ‘leaky’ expression in the absence of a driver transgene. In *Cha>GtACR2* flies, 391 µW/mm^2^ blue light induced motionless paralysis for the entire duration of light exposure (Figure 1D); an identical light level had no visible effect on control *UAS-GtACR2*/+ flies. Illumination of *Cha>eNpHR* flies with 495 µW/mm^2^ amber light failed to have any visible effect (Figure 1E); *Cha>eNpHR* animals were also placed 3 mm above an LED emitter (~1900 µW/mm^2^) which resulted in sporadic paralysis, confirming that these flies carried active eNpHR (Figure 2D-E, Video 2E). Video analysis indicated that *Cha>GtACR1* and *Cha>GtACR2* paralysis onset times were strongly dependent on light intensity, with millisecond-scale onsets occurring at powers 38 µW/mm^2^ and 391 µW/mm^2^ and above, respectively (Figure 1G, J). Recovery times were not dependent on light intensity (Figure 1H–I, K–L, Figure 2B–C).

### Profound differences between GtACR-induced cholinergic paralysis and CsChrimson-induced seizure

The GtACRs are potent inhibitors of action potential firing in several cell types (Govorunova et al. 2015; Mahn et al. 2016), however chloride conductances can have various effects on membrane potential (hyperpolarization, depolarization, shunting) depending on a cell’s chloride reversal potential (Knoflach, Hernandez, and Bertrand 2016). GtACR1 photocurrents have been observed to induce transient synaptic release at light-on, an effect attributed to chloride-mediated depolarization at the axon terminus (Mahn et al. 2016; Wiegert and Oertner 2016). We hypothesized that the GtACRs were inducing paralysis by neuronal activation rather than silencing and that an activating photochannel would thus phenocopy the GtACR-mediated paralysis. Activation of the cholinergic system with CsChrimson refuted this hypothesis, revealing that while illuminated *Cha>CsChrimson* flies underwent convulsions after light-on, followed by a tetanic pose, *Cha>GtACR* flies rapidly entered a static paralysis and retained their pre-light pose (Figure 1C–E, Figure 2F–K, Video 3). These profoundly different effects are consistent with the idea that GtACRs act to inhibit action potentials (Govorunova et al. 2015; Mahn et al. 2016).

### GtACR1 illumination silences action potentials in a nerve

Photo-actuation of GtACR1 expressed in larval abdominal nerves

(*nSyb>GtACR1*, pan-neuronal expression) produced dramatic reductions in spiking frequency, as seen in representative recordings from larval segmental nerves (Figure 1M, Figure 3). For example, 500 ms pulses of 18 µW/mm^2^ green light suppressed the spiking frequency of *nSyb>GtACR1* larval nerves by Δ∆ = –97.8% relative to the preceding activity [95CI –96.0, –98.7], *P* = 3.62 × 10-33, N_pulses_ = 22, N_flies_ = 3 (Figure 3A). At the same intensity, 30 s pulses decreased the spiking frequency by –99.8% [95CI –99.4, –100], *P* = 3.83 × 10-16, N_pulses_ = 7, N_flies_ = 3. Across several light intensities and pulse durations tested, we did not observe any decrease in the spiking frequency of control larvae during green light illumination (*nSyb-Gal4/+* and *UAS-GtACR1/+*). For instance, 500 ms pulses of 38 µW/mm^2^ green light did not alter the spiking frequency of *UAS-GtACR1/+* larval nerves (Δ∆ = –0.24% [95CI –11.6, +15.5], *P* = 0.97, N_pulses_ = 25, N_flies_ = 3, Figure 3C). These results verify that GtACRs are potent inhibitors of neuronal excitability in *Drosophila*.

**Figure 3.**
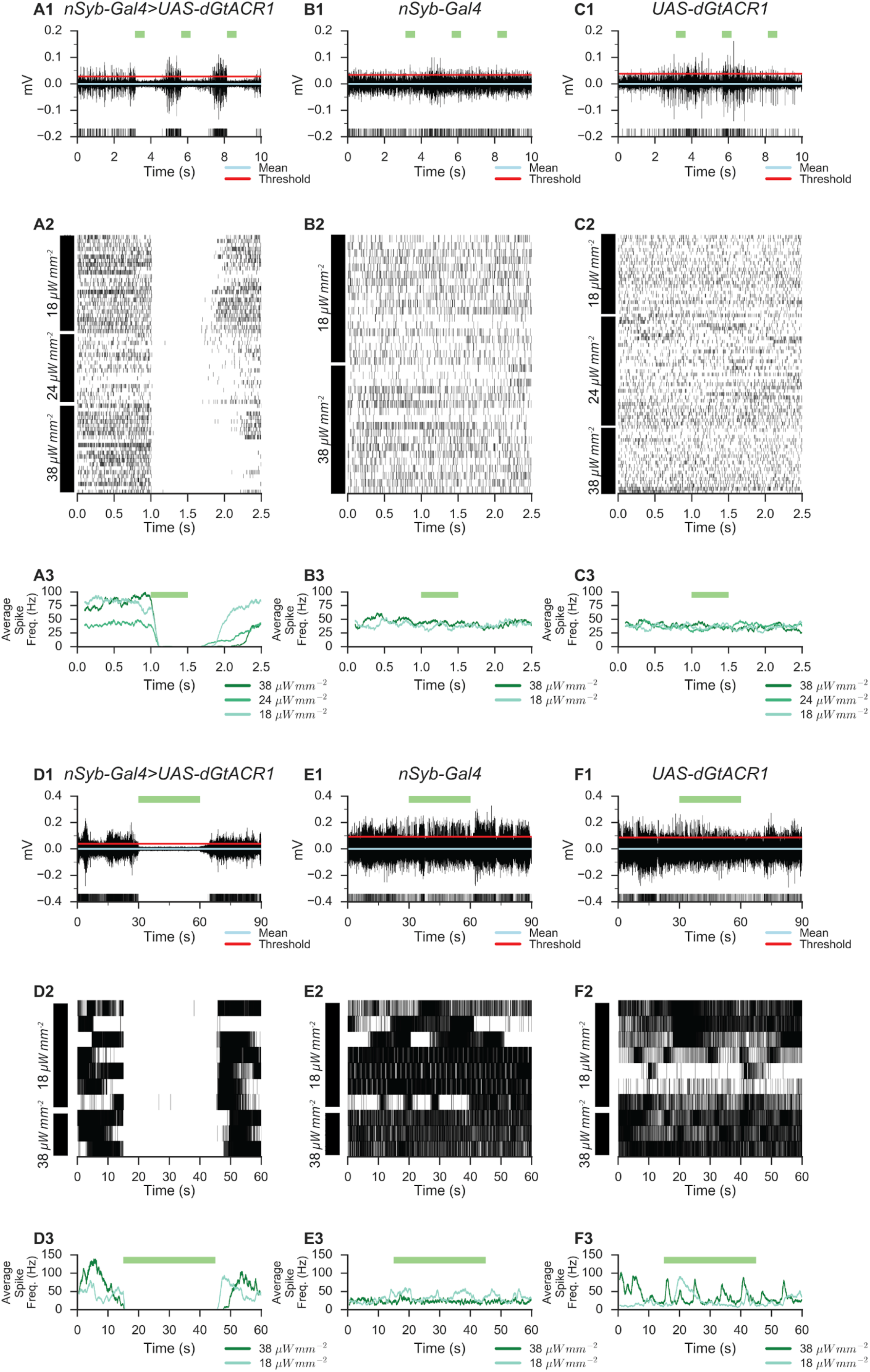
**A.** A representative trace (**A1**) from an extracellular recording from a larval abdominal 3 nerve showing the inhibitory effect of three 500 ms pulses of green light (38 µW/mm^2^) in a *nSyb>GtACR1* larva. The red line indicates the voltage threshold used to detect spikes. Rasters of spike timings (**A2**) from several light intensities (38 µW/mm^2^, 24 µW/mm^2^, and 18 µW/mm^2^) and a spike frequency plot (**A3**) indicate that the time of full activity recovery is longer with more intense light (N_pulses_ = 93, N_flies_ = 3). 500 ms pulses of 38 µW/mm2 green light nerves from *nSyb>GtACR1* larvae suppressed spiking frequency by ∆ = −98.5% [95CI −99.2, −97.6], *P* = 3.05 × 10-39, N_pulses_ = 23, N_flies_ = 3. With 500 ms pulses of 24 µW/mm^2^ green light, the spiking frequency exhibited a decrease of −99.0% [95CI −99.7, −97.7], *P* = 3.415 × 10-35, N_pulses_ = 21, N_flies_ = 3. **B.** Illumination of *nSyb-Gal4/+* control flies with green light (38 and 18 µW/mm^2^) for 500 ms had no detectable effect on the rate of identifiable spikes (N_pulses_ = 63, N_flies_ = 3). **C.** Illumination of *UAS-GtACR1/+* control flies for 500 ms with green light (38, 24, and 18 µW/mm^2^) had no effect on nerve firing (N_pulses_ = 106, N_flies_ = 3). **D.** Representative recording (D1), raster plot (D2) and frequency plot of *nSyb>GtACR1* nerves indicate that firing is almost completely suppressed during an 30 s illumination epoch. Firing recovers in <5 s at 18 µW/mm^2^ and ~10 s at 38 µW/mm^2^ (N_pulses_ = 10, N_flies_ = 3). 30 s pulses of 38 µW/ mm^2^ light completely silenced spikes, ∆ = −100% [95CI −100, −100], P = 0, N_pulses_ = 10, N_flies_ = 3. **E.** Illumination of *nSyb-Gal4/+* controls for 30 s had no effect on spiking (N_pulses_ = 10, N_flies_ = 3). **F.** Illumination of *UAS-GtACR1/+* controls for 30 s had no effect on firing (N_pulses_ = 10, N_flies_ = 3).

### Illuminating GtACRs in bursicon cells prevents wing expansion

Bursicon is a neurohormone required for developmental functions including wing expansion after eclosion; inhibition of bursicon release with expression of an inhibitory channel is frequently lethal, and in surviving flies prevents normal wing expansion (Peabody et al. 2008). Flies expressing GtACRs under the control of a *Bursicon* promoter (*Burs-Gal4*) were illuminated from 1-2 days after puparium formation (APF) until 9-10 APF before wing expansion was scored. The majority of illuminated *Burs>GtACR* flies died during development (129 out of 249), while the survivors failed to expand their wings (Figure 4A–B). Expression of an inward rectifying potassium channel (Kir2.1) (Baines et al. 2001) resulted in total wing expansion failure as previously reported (Peabody et al. 2008), but amber-illuminated *Burs>eNpHr* flies displayed normal wings.

**Figure 4.**
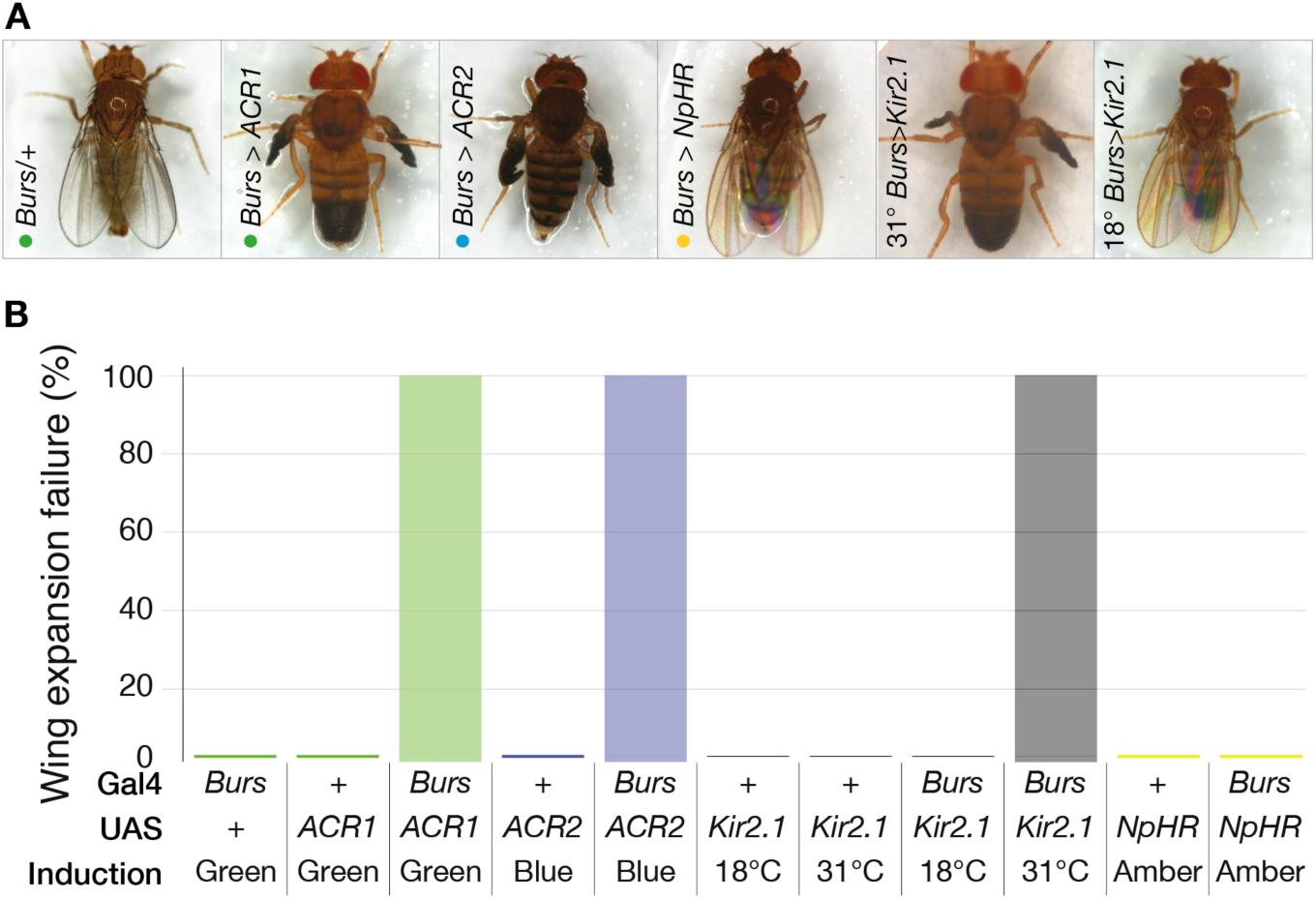
Illuminated anion channelrhodopsin expressed in *Bursicon* cells inhibit wing expansion. **A.** Illumination of flies expressing GtACRs in bursicon-releasing cells (*Burs-Gal4; UAS-GtACR1/2*) inhibited wing expansion. Flies expressing GtACR1 or GtACR2 in bursicon-releasing cells failed to expand their wings following exposure to green light at 46 μW/mm^2^ or blue light at 106 μW/mm^2^, respectively. Flies with induced Kir2.1 expression in *Burs-Gal4* cells also failed to expand their wings. Control flies were unaffected by illumination, as were and *Burs>eNpHR* flies (amber light at 106 μW/mm^2^). **B.** Illumination of flies expressing GtACRs in bursicon-releasing cells (*Burs-Gal4; UAS-GtACR1/2*) inhibited wing expansion. All flies expressing GtACR1 or GtACR2 in bursicon-releasing cells failed to expand their wings following exposure to light, controls were unaffected by light (green at 46 μW/mm^2^, blue at 106 μW/mm^2^ for GtACR1 and GtACR2 respectively). Induction of Kir2.1 at 31°C in *Burs-Gal4>UAS-Kir2.1, Tub-Gal80^ts^*flies produced a 100% failure in wing expansion. Illumination of *Burs-Gal4>UAS-eNpHR* flies with 106μW/mm^2^ amber light had no effect on wing expansion. *Burs/+* (145), *Burs-Gal4; UAS-GtACR1* (N=114), *Burs-Gal4; UAS-GtACR1* (N=135).

### Actuating GtACR1 in gustatory neurons inhibits the proboscis extension reflex

*Drosophila* extend their proboscis to sweet liquids, a response known as the proboscis extension reflex (PER) which is dependant upon the activity of neurons expressing Gustatory receptor 64f (Gr64f) (Thoma et al. 2016). Optogenetic activation with CsChrimson is sufficient to elicit PER in the absence of sugar (Klapoetke et al. 2014). When illuminated with green light, flies expressing *UAS-GtACR1* in *Gr64f-Gal4* neurons failed to exhibit PER when presented with a sucrose solution (Figure 5G−H). Exposure of the same flies to red light had essentially no effect on PER (Figure 5A−B). Flies expressing eNpHr in Gr64f showed a reduced PER relative to controls in amber light, Δ∆PER = –0.26 [95CI −0.12, –0.42]. (Figure 5A– B). Expression of Kir2.1 in Gr64f cells reduced PER dramatically, Δ∆PER = –0.96 [95CI 0.92, –1.0] (Figure 5A–B). The potent suppression of PER by illuminated GtACR1—the opposite effect to that of CsChrimson—verifies that the anion channelrhodopsin effectively inhibits activity in Gr64f cells.

**Figure 5.**
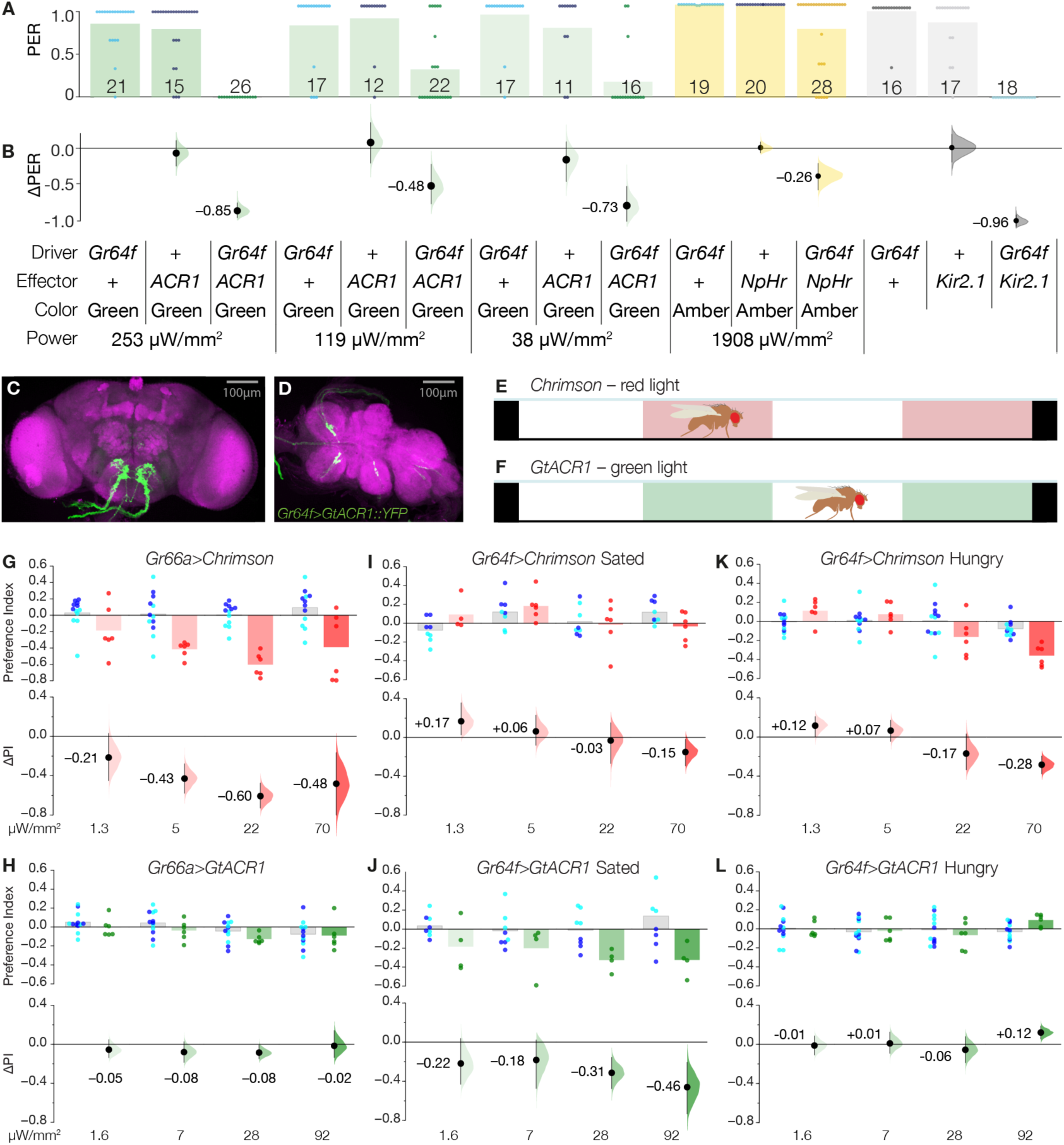
Optogenetic inactivation of taste neurons suppresses proboscis extension reflexes and flies avoid suppression of sweet taste receptors. **A.** Proboscis extension responses of transgenic flies exposed to a range of protocols aimed at silencing the Gr64f cells. Dots show individual animal response probabilities, bars represent mean response PER probabilities, numerals indicate sample sizes. Genotypes and conditions are given in key. **B.** Changes in responses (∆PER) relative to *Gr64f-Gal4/+* controls. Illumination of *Gr64f>GtACR1* flies produced effective suppression of PER at two weaker light intensities and complete suppression at the highest intensity. Flies expressing *UAS-eNpHr* in *Gr64f-Gal4* showed only a modest reduction in PER in intense amber light (1908 µW/mm^2^). Inhibition of Gr64f neurons with Kir2.1 led to a dramatic reduction in PER as compared to controls. The mean differences are as follows: *Gr64f>GtACR1* at 253 μW/mm^2^ ∆PER = −0.85 [95CI −1.02, −0.64], *P* = 3.5 × 10^−08^; 119 μW/mm^2^∆PER = −0.48 [95CI −0.21, −0.71], *P* = 1.9 × 10^−03^; 38 μW/mm^2^ ∆PER = −0.73 [95CI −0.49, −0.94], *P* = 1.7 × 10^−05^; *Gr64f>eNpHr* ∆PER = −0.26 [95CI −0.42, −0.12], *P* = 0.054 and *Gr64f>Kir2.1* ∆PER = −0.96 [95CI-1.0, −0.92], *P* = 2.7 × 10^−08^. **C–D.** The morphology of *Gr64f>GtACR1::YFP* neurites in the brain and ventral nerve cord. Magenta indicates neuropil stained with anti-DLG, green indicates YFP fluorescence. **E–F.** To examine neuronal activation preference (valence) in CsChrimson-bearing flies, a chamber was illuminated with two bands of red light. Diagram not to scale. Valence responses of GtACR1-bearing flies were examined in a chamber illuminated with bands of green light. **G.** Flies expressing CsChrimson in their bitter taste neurons (*Gr66a>CsChrimson*, red dots) were tested for their preference for red light (*Gr66a/+*, blue dots; *UAS-CsChrimson/+*, cyan dots), measured as a preference index (PI). The projector light intensities were: 1.3, 5, 22 and 70 μW/ mm^2^, left to right. Each dot indicates an experimental iteration, i.e. N = ≥4; each iteration used 15 flies. **Lower panel:** Preference relative to control animals (∆PI) was calculated as the mean difference between experimental and control PI scores: *Gr66a>CsChrimson* flies avoided activation at all intensities. At 22 μW/mm^2^ ∆PI = −0.61 [95CI –0.47, −0.73], P = 0.0009. **H.** Flies expressing *GtACR1* in their bitter taste neurons (*Gr66a>GtACR1*) had green light preferences that were similar to control animals, at all intensities (1.6, 7, 28, 92 μW/mm^2^, left to right). **I.** In a sated state, flies expressing *CsChrimson* in sweet taste neurons (*Gr64f>CsChrimson*) showed a modest attraction for 1.3 μW/mm^2^ red light, ∆PI = +0.17 [95CI 0.03, +0.37], *P* = 0.10, and a mild avoidance of 70μW/mm^2^ red light ∆PI = –0.15 [95CI +0.02, –0.29], *P* = 0.11. **J.** In a sated state, flies expressing *GtACR1* in sweet taste neurons (*Gr64f>GtACR1*) avoided green light at a variety of intensities, including strong avoidance at the highest intensity, ∆PI = –0.46 [95CI –0.19, –0.73], *P* = 0.05. **K.** In a hungry state, flies expressing *CsChrimson* in sweet taste neurons (*Gr64f>CsChrimson*) showed a similar overall response profile as the sated flies, but showed stronger avoidance at the higher intensities. **L.** Hungry flies expressing *GtACR1* in sweet tasted neurons were largely indifferent to green light, though were mildly attracted to the highest intensity ∆PI = +0.12 [95CI +0.05, +0.19], P = 0.01.

### Flies avoid GtACR1 inhibition of sweet taste receptor neurons

CsChrimson activation of bitter-sensing Gustatory receptor 66a (Gr66a) neurons was aversive to flies (Figure 5G) (Aso et al. 2014). However, *Gr66a>GtACR1* flies were indifferent to green light over a range of intensities known to have effects in other behaviors (Figure 5H), suggesting that the bitter-sensing system is quiet in the absence of a stimulus. Sated flies bearing CsChrimson in their Gr64f sweet taste cells were mildly attracted to dim red light and minimally averted by stronger red light (Figure 5I). In the hungry state, *Gr64f>CsChrimson* flies had a similar response profile, though avoided the higher intensities of red light slightly more (Figure 5K). Surprisingly, sated *Gr64f>GtACR1* flies avoided a range of green light intensities, including a strong aversion to 92 µW/mm^2^ (Figure 5J). However, hungry *Gr64f>GtACR1* flies were indifferent to green light. These results confirm that emotional responses to anion channelrhodopsin conductance in gustatory cells are distinct from responses to activator photochannels, verifying that GtACR1 is an inhibitor of neuronal activity. This experiment also demonstrates the utility of an optogenetic inhibitor to understanding how tonic activity and quiescence contribute to behavior.

### Inhibition of the mushroom body with GtACR1 suppresses short term memory performance

Synaptic transmission from the Kenyon cells of the mushroom body is required for normal olfactory short term memory (STM) (McGuire, Le, and Davis 2001; Dubnau et al. 2001; Yildizoglu et al. 2015). We hypothesized that GtACR-induced inhibition of the Kenyon cells would compromise fly performance during STM tests. We subjected *Drosophila* expressing GtACR1 under the control of a mushroom body-specific driver (*OK107-Gal4*) to aversive olfactory conditioning first under infra-red, and then green light. In all genotypes, exposing flies to infrared light during the first training cycle allowed the formation and expression of robust STM (Figure 6B–F). In *OK107>eNpHR* flies, bright amber light (1.7 mW/mm^2^) reduced STM performance, however control animals were similarly affected by the light, suggesting marginal inhibition by eNpHR action (Figure 6B). Heat treatment of flies had minor effects on controls, but produced robust STM inhibition in *OK107>shi^ts^*animals (Figure 6C). Subjecting *OK107>GtACR1* flies to green light produced a robust inhibition of performance, with negligible effects on control lines (Figure 6D–F) across a range of intensities. These data confirm that GtACR1 has similar efficacy for inhibiting central brain neurons as the current standard thermogenetic tool.

**Figure 6.**
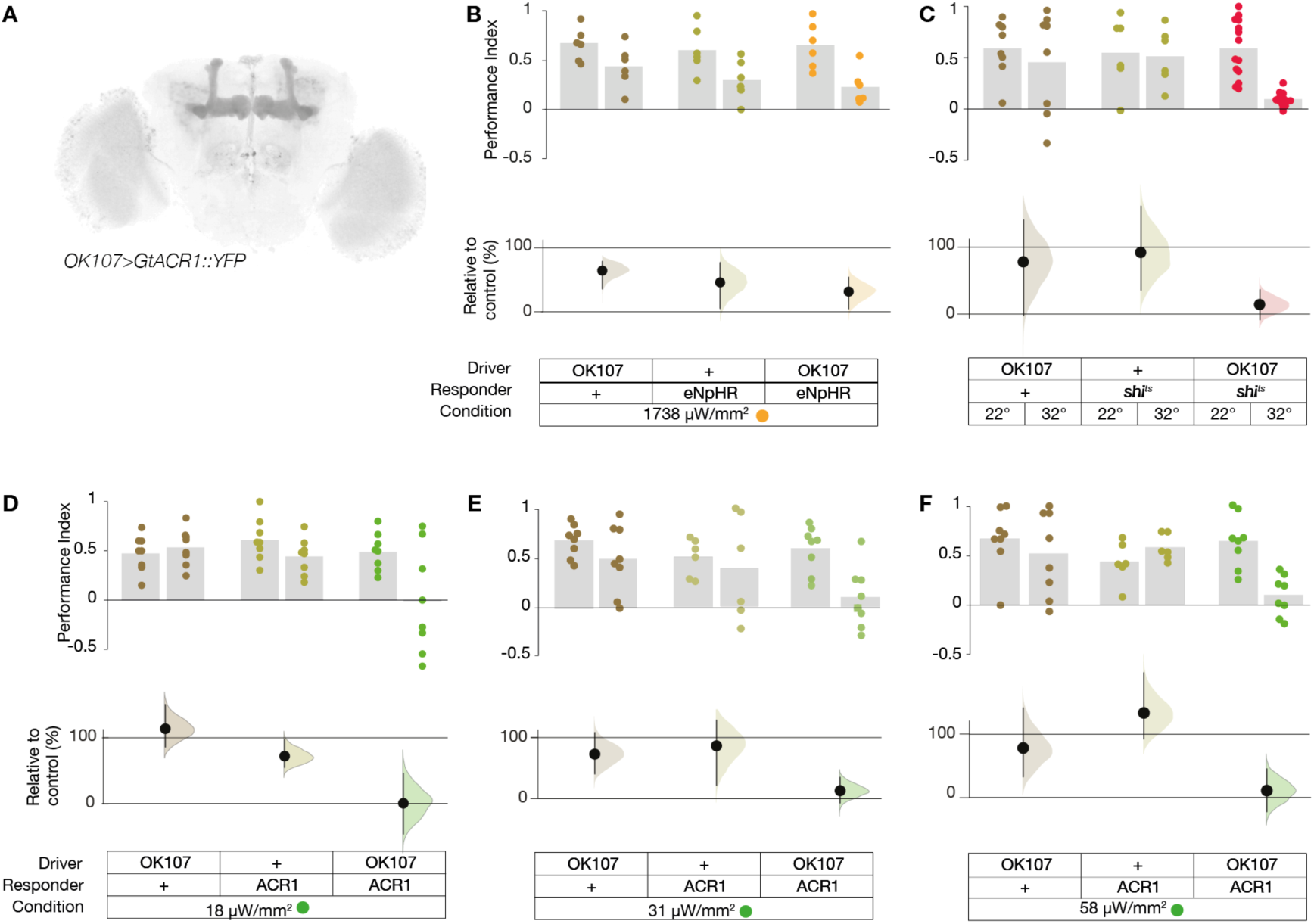
Optogenetic inactivation of Kenyon cells diminishes olfactory short-term memory. **A.** The expression pattern of *UAS-GtACR1::YFP* driven with *OK107-Gal4*. Grey indicates YFP fluorescence. **B.** Flies were trained and tested under infrared light in an initial training cycle (left columns) and then trained and retested again under optogenetic light during a second training cycle (right columns). Flies expressing eNpHR in the mushroom body cells (*OK107>eNpHR*) had a reduction in STM during strong amber illumination, ∆PI = –0.43 [95CI −0.71, −0.14], *P* = 0.005. However, this reduction was only modestly stronger than control animals, suggesting the effect was primarily due to the light alone. Genotype key is at the bottom, colored dots represent the average of two half PIs from 6 animals each, N = 8 experiments as indicated by the dots. **Lower axis:** The memory effects of illumination as a percentage of the same animals’ scores under infrared light. Error bars are confidence intervals of the mean; curve is the bootstrap distribution of the mean. **C.** Inhibition of *OK107* cells with *UAS-shi*^ts^ at 32°C led to an almost complete block of STM, ∆PI = – 0.5 [95CI −0.67, −0.33], *P* = 1.4 × 10^−05^. A similar heat treatment of controls produced only trivial effects. **D.** Illumination with green light either slightly increased (*OK107*/+) or decreased (*UAS-GtACR1*/+) STM performance in control animals. However, green light illumination at 18 μW/mm^2^ reduced STM performance in flies expressing *OK107-Gal4>UAS-GtACR1*. ∆PI = –0.50 [95CI −0.97, −0.1], *P* = 0.03. **E.** Illumination of *OK107>GtACR1* flies with 31 μW/mm^2^ green light also inhibited the expression of STM ∆PI = –0.5 [95CI –0.33, –0.66], *P* < 0.001. **F.** Illumination of *OK107>GtACR1* flies with 58 μW/mm^2^ light inhibited conditioned avoidance ∆PI = –0.55 [95CI –0.31, –0.76], *P* = 0.003.

### The GtACRs are minimally toxic

We sought to estimate the toxicity of GtACR expression and/or activation. The expression of GtACRs in *corazonin* neurons had little to no effect on neuronal morphology, even after 6 days of continual light exposure (Figure 7A–I). Expression of GtACRs in fly ommatidia with light exposure throughout metamorphosis had no effect on eye morphology (Figure 7G-H). Moreover, broad expression of GtACRs in cholinergic neurons with intermittent light exposure did not dramatically shorten lifespans relative to controls (Figure 7J–K).

**Figure 7.**
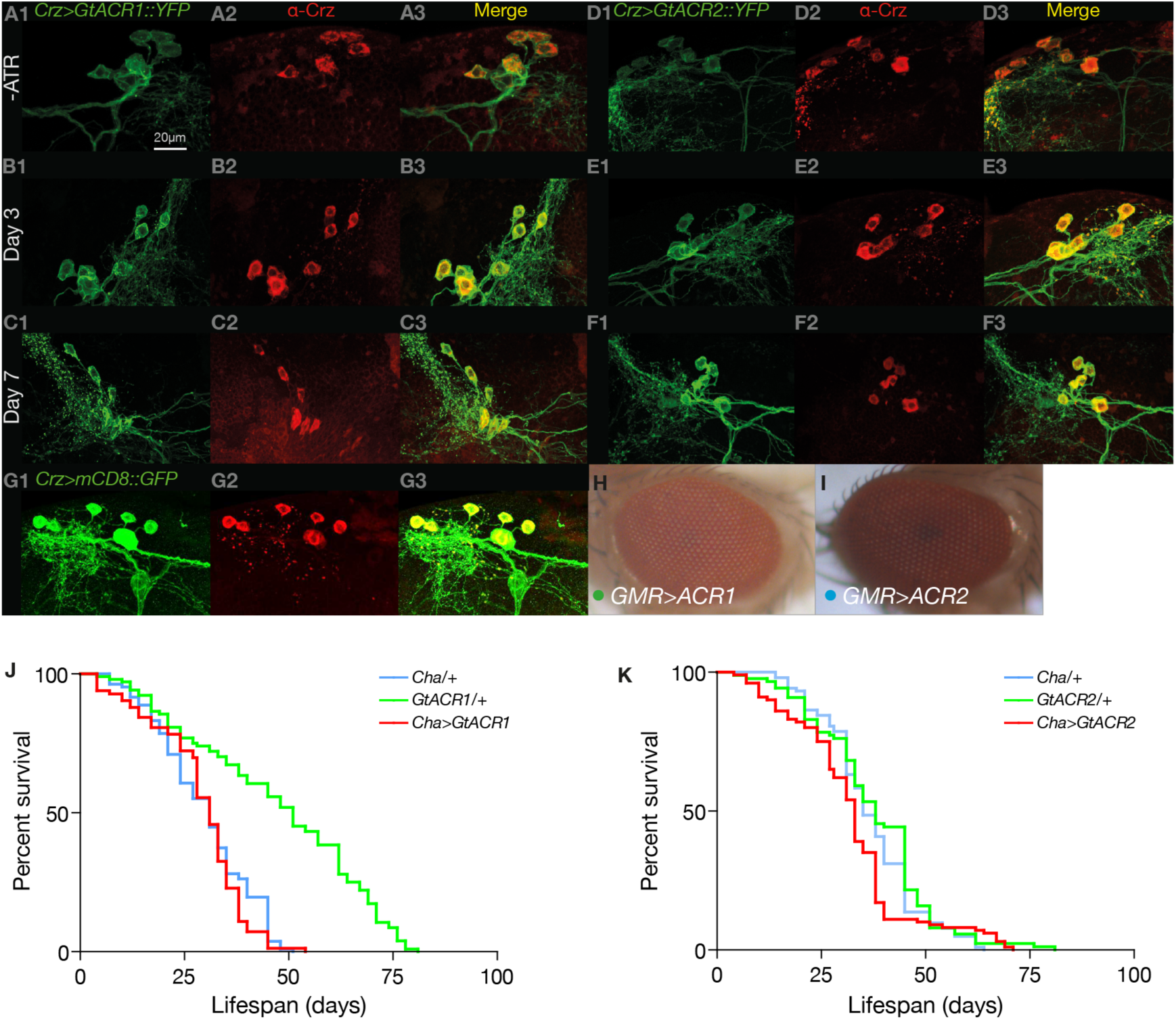
GtACRs have minimal toxicity. **A1.** Central brain cells were observed to express yellow fluorescent protein-tagged GtACR1 (green, endogenous fluorescence) in *Crz>GtACR1* fly brains prior to all-*trans*-retinal (ATR) feeding. **A2–3.** Antibody staining for Crz protein (α-Crz, red) confirmed GtACR1-expressing cells were Crz^+^. **B1–3.** Crz+ cell morphology was normal in *Crz>GtACR1* flies after ATR feeding and one day of illumination with 19 μW/mm^2^ green light. **C1–3.** Crz+ cell morphology was normal in *Crz>GtACR1* flies after ATR feeding, 6 days of illumination. **D–F.** Flies expressing GtACR2::YFP in Crz cells had normal morphology after 6 days of GtACR2 activation with 25 μW/mm^2^ blue light. **G.** The morphology of cells carrying a fluorescent protein (mCD8::GFP) but no photochannel. The mCD8::GFP was visualized with anti-GFP antibody, thus appears brighter. **H–I.** Flies expressing either GtACR1 or GtACR2 with the *GMR-Gal4* driver had normal eye morphology after being illuminated during metamorphosis. **J.** The lifespans of flies bearing *Cha>GtACR1* transgenic expression (median = 31 days [95CI 28, 33]) were similar to *Cha-Gal4/+* controls (median = 35 days [95CI 33, 40]), and shorter than the lifespans of *GtACR1/+* animals (median = 51 days [95CI 45, 57]). All flies were subjected to intermittent *ad hoc* light exposure. **K.** The lifespans of flies with *Cha>GtACR2* transgenic expression (median = 31 days [95CI 28, 33]) were similar to *Cha-Gal4/+* (median = 31 days [95CI 27, 33]) and *GtACR2/+* (median = 38 days [95CI 33, 45]) controls.

## Discussion

The *Guillardia theta* anion-conducting channelrhodopsins are highly effective tools for the optogenetic inhibition of behavioral circuits in freely-moving animals. The present data reveal that the GtACRs have comparable potency with widely used transgenic and thermogenetic neuronal inhibitors, with temporal precision similar to that of optogenetic activators. For example, the GtACRs have comparable efficacy to *shi*^*ts*^ (Figure 6), but are much faster (Figure 1G)(Kitamoto 2001). We have shown that the GtACRs are more potent than the chloride pump eNpHR, which requires very strong light intensities (Wu et al. 2014; Inada et al. 2011). Indeed, GtACR1 is so effective that intensities as low as 119 µW/mm^2^ can elicit immobilization in a fraction of *UAS-GtACR1*/+ control flies, necessitating calibration of light power. Millisecond-scale, complete paralysis was seen in *Cha>GtACR1* flies at power as low as 38 µW/mm^2^ (Figure 1G), while no immobilization was observed in *UAS-GtACR1*/+ with light as strong as 92 µW/mm^2^ (Figure 2J). This makes 38–92 µW/mm^2^ the recommended light range for adult *Drosophila* experiments with GtACR1. For GtACR2, we recommend ~391 µW/mm^2^, a blue light power that had no visible effect on control *UAS-GtACR2/+* animals while eliciting fast and complete cholinergic paralysis in *Cha>GtACR2* flies (Figure 1J). In practical terms, this means GtACR1 can be actuated with a projector, while GtACR2 requires LED arrays for most freely moving adult *Drosophila* applications. The GtACRs represent a dramatic improvement over existing methods, orders of magnitude faster than the thermogenetic tools and greatly more potent than eNpHR. The GtACRs are an important addition to the optogenetic toolkit and make neural activity necessity tests as accessible as sufficiency tests are now. **⁂**

## Methods

### Transgenes, fly strains and rearing conditions

*Drosophila melanogaster* flies were used in all experiments. *ChAT-Gal4.7.4* (BL 6798) (Salvaterra and Kitamoto 2001), *Burs-Gal4* (BL 40972)(Peabody et al. 2008), *20x-UAS-eNpHr3* (BL 36350) (Petersen and Stowers 2011), *20x-UAS-CsChrimson* (BL 55134) (Klapoetke et al. 2014), *Gr64f-Gal4* (BL 57699) (Weiss et al. 2011), were obtained from the Bloomington Drosophila Stock Center. *OK107Gal4* (DGRC 106098) (Connolly et al. 1996) was obtained from the Kyoto Stock center. GMR-Gal4 flies was a gift from Gerry Rubin (Freeman 1996). *Crz-Gal4* was a gift from Jay Park (Choi, Lee, and Park 2006). *w^1118^* flies were used as wild-type controls in all experiments. To generate *UAS-GtACR1* and *UAS-GtACR2* transgenic lines, *Drosophila*-codon-optimized sequences of *GtACR1* and *GtACR2 (Govorunova et al. 2015)* were synthesized *de novo* (GenScript Pvt. Ltd) as EYFP fusions and subcloned into *pJFRC7-20XUAS-IVS-mCD8::GFP*, acquired from Addgene (addgene.org, plasmid #26220) (Pfeiffer et al. 2010) by replacing the *mCD8:GFP* fragment with *GtACR1-EYFP* or *GtACR2-EYFP* fragments, followed by sequence verification (GenScript). Constructs were injected into an *attP2* insertion site on the third chromosome, and the transgenic progeny were balanced (BestGene Inc.). Expression of the GtACR constructs was verified by YFP fluorescence. For adult fly experiments, 3-4 days old male flies were fed with 1mM all-*trans*retinal (Sigma) for 2-3 days at 25°C in the dark. A stock solution of all-*trans*-retinal was prepared in 95% ethanol (w/v) and mixed in with warm and liquefied fly food. Each vial was covered with aluminium foil and placed in the dark.

### Optogenetic photoactivation

Green, blue, red and amber LEDs (LUXEON Rebel LED on a

SinkPAD-II 10mm Square Base available from www.luxeonstar.com; green SP-05-G4, peak emission 525 nm; blue SP-05-B4, peak 460nm; red-orange SP-05-E4, peak 617 nm and pc-amber SP-05-A5, peak 591 nm), or an LED micro-projector (Optoma ML750), were placed near behavioral arenas to provide an illumination source. LEDs were powered at maximum brightness by a 700mA BuckPuck driver and illumination intensity was controlled by varying the distance between the source and subject. The PC-controlled LED micro-projector was used to project images consisting of entirely red, green or blue backgrounds onto the arena. The projector uses a Digital Light Processing (DLP, Texas Instruments) micromirror device with a refresh rate of 60 Hz. For a consistent environment, the illuminatory and behavioral monitoring system was placed inside the incubator and maintained at 25°C throughout the experiment.

### Video acquisition

Videos were captured at 30 frames per second using an AVT Guppy F-046B CCD camera (Stemmer Imaging, UK), equipped with a 12 mm CCTV-type lens and connected to a computer via an IEEE 1394 cable. Experiments were conducted under infrared (IR) light; an IR longpass filter (Edmund Optics, Singapore) was used to reduce detection of the micro-projector/LED light. Custom-built image acquisition software CRITTA, written in LabView (National Instruments, USA), was used to track the movements of animals in the behavioral arenas.

### Light intensity measurements

The light intensities of the projector and LED illumination were measured for all wavelengths for every configuration used in the experiments: number of LEDs, distance from chamber, type of lens and projector DLP intensity values. A thermal power sensor (Thorlabs S310C) connected to a power and energy meter console (Thorlabs PM100D) was used to measure power in a dark room. The meter was zeroed before each set of measurements and a cardboard shield with a 20 mm diameter cutout was used to ensure light only struck the sensor’s absorbent surface.

### Falling assay

Fly climbing/locomotion performance was monitored in a custom acrylic arena. Four ATR treated flies, cooled on ice were transferred into each arena. The climbing was performed in a rectangular arena (70 × 11 mm), 4 such arenas were cut into a 1.5mm thick transparent acrylic sheet that was incorporated into a transparent acrylic ‘sandwich’. Climbing behavior was recorded under IR backlighting. The arena sandwich was illuminated from the front by DLP projector. Flies were allowed to freely explore the arena during the test session. The behavior of flies illuminated with IR, green, blue and red light was recorded. For higher intensity, the projector was placed closer to the behavioral arena. For the duration of the test, flies were individually tracked using a monochrome camera connected to CRITTA tracking software which also controlled the timing, hue and intensity of the illumination by driving the projected image. As well as recording the vertical position of each fly, the software divided the chamber into two zones, the lower 5.5% and the upper 94.5% and the number of flies occupying each zone was logged. Fully paralysed flies fell to the bottom of the chamber, occupied the lower zone and were scored as fallen.

### Walking paralysis

A Guppy F-046B camera was used to record fly behavior at 25 frames per second in stadium-shaped arenas (55 × 4 × 1.5 mm); fifteen such arenas were cut from acrylic. *Cha>CsChrimson*, *Cha>GtACR1, Cha>GtACR2*, and control flies were illuminated with a DLP projector. CRITTA and Python scripts were used to analyze walking speeds and generate the Muybridge series.

### High frame-rate video analysis of walking paralysis

High frame video was recorded at 1000 frames per second using a Photron FASTCAM MC2 Camera, was conducted to quantify the onset of inhibition and recovery with high temporal accuracy. For each genotype studied, 4 flies were anesthetized on ice and loaded into 4 separate clear acrylic chambers measuring 9 × 9 × 1.5mm and covered by a microscope cover slip. For GtACR recordings the chamber was front-illuminated by a red-filtered fiber optic light source and for eNpHR, which is known to be sensitive to red light, IR backlighting was used. Before recording fly locomotion was elicited by shaking the chamber. Red, green, blue or amber LEDs, set at varying distances from 3mm to 330mm from the subject, were turned on manually when the fly was freely walking and turned off several seconds after paralysis occurred. The video buffer was limited to 16 seconds, and the illumination duration was typically 2 s, though prolonged to a maximum of 30 s if no effect was seen. Videos were analysed offline by visual inspection to determine the frame where flies were first immobilized, defined as when forward locomotion is abated (though some movement may still occur due to momentum or gravity); the frame containing the first sign of recovery, e.g. the movement of a leg; and the first frame where the fly has recovered, is fully mobile and begins walking away. In addition, the frames where the light is turned on and off were marked and then the following timings were calculated: time from light onset to immobilization, time from light-off to first recovery and time from light-off to re-mobilization. Pixel difference data was analyzed with custom scripts in LabVIEW and Python.

### Wing expansion assay

For wing expansion experiments, larvae were fed with 200µM all-*trans*-retinal (ATR) throughout larval development. At 1-2 days after puparium formation (APF), pupae were transferred to green, blue or red light conditions at 25°C. Flies remained under this light until 9-10 days APF, when wing expansion was scored under a dissection microscope. For Kir2.1, all flies were raised at 18°C until 1-2 days APF, pupae were transferred to either 18°C or 31°C for *Tub-Gal80^ts^*de-repression of Kir2.1 expression. Wing expansion was scored under a dissection microscope until 9-10 days APF.

### Olfactory short term memory

Aversive olfactory conditioning of *Drosophila* was performed in custom-built chambers modified from a previously described single-fly olfactory trainer apparatus (Claridge-Chang et al. 2009). To allow light penetration and video monitoring of multiple flies, windows were cut into the top and bottom of each chamber. The floor and ceiling of each chamber was a glass slide printed with transparent indium tin oxide (ITO) electrode boards (Visiontek UK) (Vogt et al. 2014). Each side of the ITO board was sealed by a gasketed lid that formed a seal around the gap between the ITO board and the chamber wall. The internal behavioral arena measured 50 mm long, 5 mm wide and 1.3 mm high. Mirrors were aligned at a 45° angle and placed into holders on top of each chamber. Facilitated by carrier air, odors entered each end of the chamber via two entry pipes and left the chamber through two vents located in the middle of the chamber. Flies were conditioned using electric shocks (12 electric shocks at 60 V) that were through the circuit boards. At the start of each conditioning experiment the flies were iced in darkness and loaded into the chambers, 6 flies per chamber. Experiments were performed with four chambers simultaneously that were plugged into a rack in a 2 × 2 manner. Throughout the conditioning cycle, *Drosophila* flies were entrained to either avoid 3-octanol (OCT) or 4-methylcyclohexanol (MCH). Conditioning performance was tested by exposing one half of the chamber to the punished odor and the other half to the unpunished odor. A performance index (PI) was calculated by counting flies in individual video frames over the last 30s of assessment, using a formula described previously (Quinn, Harris, and Benzer 1974). After the initial conditioning cycle, the same organisms were entrained to the same odor again by using an identical protocol except that green (**λ** 525 nm) or amber light (**λ** 591 nm) was turned on during memory retrieval. In *shi*^*ts*^ experiments, all flies were initially conditioned at 22°C and subsequently incubated for 30 minutes at 32°C to inactivate endocytosis. After the incubation period, the flies were conditioned against the same odor a second time at 22°C. Finally, the fly performance between the first and second conditioning cycle was compared.

### Optogenetic valence assay

Each arena had a 55 × 4 mm stadium layout; 15 such arenas were cut from 1.5 mm thick transparent acrylic. During an experiment, all arenas were covered with a transparent acrylic lid. Ice-anesthetized flies were loaded into each chamber in dark and the whole arena was kept under infrared light at 25°C for 2-3 minutes before starting the assay; behavior was recorded under IR lighting. The arena was illuminated from the top with visible light from a mini-projector (Optoma ML750). For CsChrimson experiments, four red light intensities were used; for *ACR* experiments, green light four green light intensities was used. The coloured light intensity was varied by changing the level of the respective digital color components of the projection. For each batch of experiments, two light-test sessions were conducted, separated by 10s. For the first test session, the arenas were illuminated for 60 seconds with stripes consisting of two light and two dark zones, all equally sized. For the second 60 second test session the locations of the light and dark zones were reversed. For the duration of the test, the positions of the flies were individually tracked using a Guppy F-046B camera with an IR bandpass filter, connected to CRITTA tracking software which also controlled the timing, hue and intensity of the illumination by driving the projected image, as well as counting the number of flies in each zone. Fly preference for light was calculated as a preference index by subtracting the total number of flies in the dark from the total number of flies in the light, and dividing this number by the total number of flies in the experiment, for each of the 4 light intensities.

### Proboscis extension reflex assay

In all experiments, 4-5 days old, 30 hr wet-starved (0.5% agarose) male flies were used. Each fly was glued by its back onto a glass slide with nail polish, then placed in a vial with a water-soaked tissue to recover for 1-2 hours. The PER response was tested by manually presenting a drop of 1 M sucrose solution (Sigma) to the forelegs for up to 5 sec, using a 1 ml syringe, in the presence of red or green or amber light. When the fly extended its proboscis, the syringe was immediately withdrawn to prevent drinking. GtACR flies were tested with red, green and blue; eNpHR was tested with high intensity amber. Flies were given water before the experiment and after each light change. Light color was randomized for each fly. Each presentation was recorded manually. Fly responses were counted offline manually. Each fly was tested 3 times, and responses were counted as either 0 or 1 (for the absence and presence of PER respectively): the mean outcome of 3 presentations was denoted as fly performance. Δ∆PER values for each condition were calculated by subtracting PER performance of the driver controls from the PER response of responder control and experimental animals.

### Survival analysis and toxicity assays

We performed survival assays of flies expressing GtACRs in all cholinergic neurons using *Cha-Gal4* after treatment with ATR for 4 days. Animals were briefly exposed to light at day 4, 7 and 12 after eclosure. Throughout the assay, *Drosophila* were transferred into new food vials every third day and any deaths were recorded; longevity was monitored until all flies were dead. To examine GtACR cellular toxicity, we expressed GtACRs in retinal cells with the Glass Multiple Reporter *GMR-Gal4* driver and examined the offspring for rough-eye phenotypes (Van Vactor et al. 1991). Larvae were kept in the dark on ATR food throughout development. Pupae were exposed to light from 2-3 days after pupal formation until eclosion, and eyes were examined 5 days after eclosion. To monitor toxicity in central brain cells, we expressed GtACR::YFP in *Corazonin (Crz)* cells with *Crz-Gal4* which expresses in 6-8 neurons (Choi, Lee, and Park 2006). After eclosion, flies were transferred to the ATR treated food for 2-3 days. Brains were dissected at three stages: before ATR treatment, 1 day after ATR treatment with light exposure and 6 days after ATR with light exposure.

### Immunohistochemistry

Adult brains were dissected in PBS and fixed in 4% paraformaldehyde for 20 min at room temperature. Samples were washed three times in PBT (phosphate buffered saline with 1% Triton X-100 at pH 7.2) and blocked with 5% normal goat serum for one hour. Samples were then incubated with primary antibodies overnight at 4°C. After three additional washes with PBT, samples were incubated with a secondary antibody overnight at 4°C. Stained brains were mounted in Vectashield (Vector Laboratories, Burlingame, CA, USA) and recorded with confocal fluorescence laser scanning microscopy (Zeiss). Anti-Crz antibody (1:1000) and goat anti-rabbit-Alexa Fluor 568 (A-11011, Molecular Probes, 1:200 dilution) were used. GtACR1::YFP and GtACR2::YFP were visualized without antibody staining. Control mCD8::GFP expression was visualised using anti-GFP (ab13970) and anti-chicken-Alexa Fluor 488 antibody staining.

### Electrophysiology

Third instar larvae were dissected in HL3 solution modified from a previous recipe as follows: 110 mM NaCl, 5 mM KCl, 5 mM HEP-ES, 10 mM NaHCO_3_, 5 mM trehalose, 30 mM sucrose, 1.5 mM CaCl_2_, 4 mM MgCl_2_ (Verstreken et al. 2003). An individual abdominal nerve (A 3/4) was drawn into a fire-polished glass suction electrode as described previously (Tracey et al. 2003). Extracellular recordings of action potentials from both sensory and motor neurons were performed using a DAM-50 Differential Amplifier (World Precision Instruments) in the AC mode at 1000× gain and bandpass filtered at 100 Hz – 1.5 kHz. Optogenetic silencing of the segmental nerve was achieved using 500 ms and 30 s pulses of light from a green LED triggered from the Clampex software (Molecular Devices) during recordings. Spiking activity occurred in bouts separated by intervals of quiescence; after recording, if no spiking was observed prior to light onset, that epoch’s data were excluded from the analysis. Spike detection was performed using a window discriminator (Meliza and Margoliash 2012). A spike was defined as an upward signal that peaked within 50 ms, and had an amplitude threshold of 2.58 standard deviations from the mean amplitude. The spiking frequency of each nerve was calculated with a rolling window of 100 ms and 500 ms, for 500 ms and 30 s illumination pulses respectively.

### Statistics and analysis

Estimation statistical methods were used to analyze and interpret quantitative data (Altman et al. 2000; Claridge-Chang and Assam 2016; Cumming 2012). For each olfactory STM experiment, the mean difference in PI for green light (relative to IR light) was computed (Δ∆PI). For each PER experiment, the mean difference in PER scores relative to driver controls was computed (Δ∆PER). Data were presented as mean difference contrast plots (Gardner and Altman 1986; Cumming 2012). Bootstrap methods (Efron 1979) were used to calculate 95% confidence intervals for the mean difference between control and experimental groups. Confidence intervals were bias-corrected and accelerated (DiCiccio and Efron 1996), and were displayed with the bootstrap distribution of the mean; resampling was performed 2,000 times. All reported *P* values are the results of two-tailed Student *t*-tests. Data analysis was performed and visualized in LabVIEW, in Matlab, and in Python using Jupyter and the scikits-bootstrap, seaborn, and SciPy packages.

### Author Contributions

*Conceptualization*: FM and ACC; *Methodology*: FM, JCS and ACC; *Software*: JCS (CRITTA, Labview) and JH (Python); *Investigation*: FM (transgene design, genetics, paralysis, wing expansion, toxicity in eyes, proboscis extension reflex, valence, neuroanatomy), SO (learning and survival), JCS (paralysis), JYC (brain dissection, immunohistochemistry, microscopy) and KC (proboscis extension reflex), TWK (extracellular recordings); *Resources*: JCS (instrumentation); *Data Analysis*: JH (valence, electrophysiology), KC (PER), JCS (paralysis, falling), SO (STM) and FM (paralysis, valence, anatomy); *Writing – Original Draft*: FM and ACC; *Writing – Revision*: FM, SO, JCS and ACC; *Visualization*: FM, JH, JCS, SO, KC and ACC; *Supervision*: ACC; *Project Administration*: ACC; *Funding Acquisition*: ACC.

## Acknowledgements

We thank John Spudich for providing the GtACR1 and GtACR2 sequences and plasmids. We thank Jan Adrianus Veenstra for the anti-Crz antibody. We thank Sherry Aw for loan of the high-speed camera. We thank George Augustine for comments on the manuscript. We thank Lucy Robinson of Insight Editing London for editing of the manuscript. FM, SO, JYC and ACC were supported by grant MOE-2013-T2-2-054 from the Ministry of Education; JCS and ACC were supported by grants 1231AFG030 and 1431AFG120 from the A*STAR Joint Council Office. JH was supported by the A*STAR Graduate Academy. The authors were supported by a Biomedical Research Council block grant to the Institute of Molecular and Cell Biology. FM, SO, KC and ACC received support from Duke-NUS Medical School, including the Integrated Biology and Medicine doctoral program (KC).

## Video Legends

**Video 1. GtACR flies fall from a vertical surface when illuminated**

Flies expressing one of three optogenetic inhibitors in their cholinergic neurons (*Cha-Gal4>UAS-GtACR1, Cha-Gal4>UAS-GtACR2* and *Cha-Gal4>UAS-eNpHR*) were illuminated with light from a projector. *Cha>GtACR1* and *Cha>GtACR2* flies fell from the vertical acrylic surface upon exposure to green or blue light respectively, and were immobilized. *Cha>GtACR2* flies retained some motor activity while illuminated with blue light. *Cha>eNpHR* flies did not fall upon exposure to red light and remained mobile.

**Video 2. GtACR flies are immobilized by illumination**

**A.** Green light at 38 μW/mm^2^ rendered a *Cha>GtACR1* fly immobile, though it regained some motor control during illumination. Green dot indicates when light was turned on.

**B.** Illumination of a *GtACR1/+* fly with 38 μW/mm^2^ green light had no effect.

**C.** A *Cha>GtACR2* fly was rendered completely paralyzed by illumination with 391 μW/mm^2^ blue light. Blue dot indicates when light was turned on.

**D.** A *GtACR2/+* fly was unaffected by illumination with 391 μW/mm^2^ blue light.

**E.** While positioned 3 mm above an amber LED (approximately 1.9 mW/mm^2^), a *Cha>eNpHR* fly retained mobility, though it was paralyzed transiently when passing directly above the emitter. Light was on throughout this recording.

**F.** A *Cha>eNpHR* fly was unaffected by amber illumination at 495 μW/mm^2^. Amber dot indicates when light was on.

**Video 3. *Cha>GtACR* and *Cha>CsChrimson* flies have distinct responses to actuation.**

*Cha>GtACR* flies adopt a static pose during illumination (indicated by colored dots), but Cha>Chrimson flies have active seizures and adopt a tetanic pose with extended wings. Control animals were unaffected by projector light (green 92 μW/mm^2^; blue 67 μW/mm^2^; red 70 μW/ mm^2^).

